# Leveraging a Vision-Language Model with Natural Text Supervision for MRI Retrieval, Captioning, Classification, and Visual Question Answering

**DOI:** 10.1101/2025.02.15.638446

**Authors:** Nikhil J. Dhinagar, Sophia I. Thomopoulos, Paul M. Thompson

## Abstract

Large multimodal models are now extensively used worldwide, with the most powerful ones trained on massive, general-purpose datasets. Despite their rapid deployment, concerns persist regarding the quality and domain relevance of the training data, especially in radiology, medical research, and neuroscience. Additionally, healthcare data privacy is paramount when querying models trained on medical data, as is transparency regarding service hosting and data storage. So far, most deep learning algorithms in radiologic research are designed to perform a specific task (e.g., diagnostic classification) and cannot be prompted to perform multiple tasks using natural language. In this work, we introduce a framework based on vector retrieval and contrastive learning to efficiently learn visual brain MRI concepts via natural language supervision. We show how the method learns to identify factors that affect the brain in Alzheimer’s disease (AD) via joint embedding and natural language supervision. First, we pre-train separate text and image encoders using self-supervised learning, and jointly fine-tune these encoders to develop a shared embedding space. We train our model to perform multiple tasks, including MRI retrieval, MRI captioning, and MRI classification. We show its versatility by developing a retrieval and re-ranking mechanism along with a transformer decoder for visual question answering.

**Clinical Relevance:** By learning a cross-modal embedding of radiologic features and text, our approach can learn to perform diagnostic and prognostic assessments in AD research as well as to assist practicing clinicians. Integrating medical imaging with clinical descriptions and text prompts, we aim to provide a general, versatile tool for detecting radiologic features described by text, offering a new approach to radiologic research.

## I. Introduction

Alzheimer’s disease (AD) is a progressive neurodegenerative disease and the leading cause of dementia worldwide. In the United States, it is estimated that over 6 million individuals aged 65 and older are living with AD [1]. Globally, the burden of AD is substantial and is projected to increase dramatically in the coming decades as populations age [2].

Accurate diagnosis and prognosis of AD are critical for effective disease management. Current diagnostic criteria, as established by the National Institute on Aging (NIA), along with the Alzheimer’s Association, rely on a combination of clinical assessments and biomarker evidence. Imaging techniques such as positron emission tomography (PET) along with structural imaging methods such as magnetic resonance imaging (MRI), play a role in this diagnostic process [3]. In particular, T1-weighted brain MRI is widely used in both clinical practice and research settings as it maps and quantifies brain anatomy in detail, making it a valuable tool for assessing atrophy patterns associated with normal aging and AD [4].

Beyond imaging, multimodal data—including clinical evaluations, cognitive tests, and other biomarkers—are collected to capture the full complexity of AD pathology [5]. Integrating diverse data sources holds great promise for improving diagnostic accuracy and understanding factors that affect disease progression. Recent advances in artificial intelligence (AI), particularly in machine learning and deep learning, have further enabled researchers to analyze large, heterogeneous datasets. These AI-driven methods include convolutional neural networks, vision transformers [6] and latent diffusion models [7] that can be trained to identify subtle imaging biomarkers for disease classification or prognosis, as well as image segmentation and labeling [8]. Extensive benchmarking [9] [10] has led to powerful methods for disease detection. Even so, AI algorithms in radiology are still typically trained to perform a single task (e.g., disease classification) and cannot usually be trained to perform multiple tasks. Here we address this problem by training a vision-language model to perform multiple radiologic tasks (image retrieval, captioning, and classification) via natural language supervision.

In computer vision, vision-language models have evolved from systems that classify images into a fixed set of categories to models that leverage natural language supervision for broader understanding. In the 2010s, models were typically trained to predict predetermined object categories. OpenAI’s CLIP model [11] shifted this paradigm by training a high-dimensional joint text-image embedding. The CLIP model was created by applying contrastive learning to 400 million image–text pairs, thereby achieving strong zero-shot classification without task-specific tuning. Subsequent developments include the Large Language and Vision Assistant (LLaVA) [12], which integrates a vision encoder with a large language model for end-to-end multimodal understanding. More recent advancements include Google’s PaliGemma2 [13], built on the Gemma2 language model, along with specialized models such as BioMedGPT [14] for biomedical tasks. In the medical imaging domain, MedBLIP [15] —a BLIP-2-based architecture—extends 2D models to 3D data, while NeuroBERT [16] demonstrates the potential of text-vision approaches for detecting abnormalities in brain MRI scans.

Most large-scale vision-language models lack the capability to handle 3D medical imaging data natively. Using proprietary models raises concerns about training data quality, privacy issues and whether the training data is sufficiently relevant to be useful in specific domains such as radiology and clinical neuroscience. As well as imaging inputs, often demographic variables such as the patient’s age and sex [17] are important covariates for gauging AD pathology and disease-related atrophy, but they are often not readily integrated into diagnostic models that learn from images.

In this paper, we propose a novel vision-language framework as shown in **Figure 1**, designed to augment brain MRI analysis by integrating T1-weighted imaging data with text descriptions of subject-specific information, such as age, sex, and diagnosis. We evaluated our cross-modal model on multiple tasks - MRI retrieval, MRI captioning, and MRI classification. We illustrate the capability of our joint text-image vector embeddings with a retrieval and re-ranking mechanism using a text decoder for visual question answering. We conduct several ablation experiments to show the contributions of different components in our proposed framework. By training a deep learning model to learn connections between structural imaging and text-based data, we offer a comprehensive framework that could advance radiologic research, offering, in principle, a new approach to discover factors that affect disease progression.

**Figure 1.**
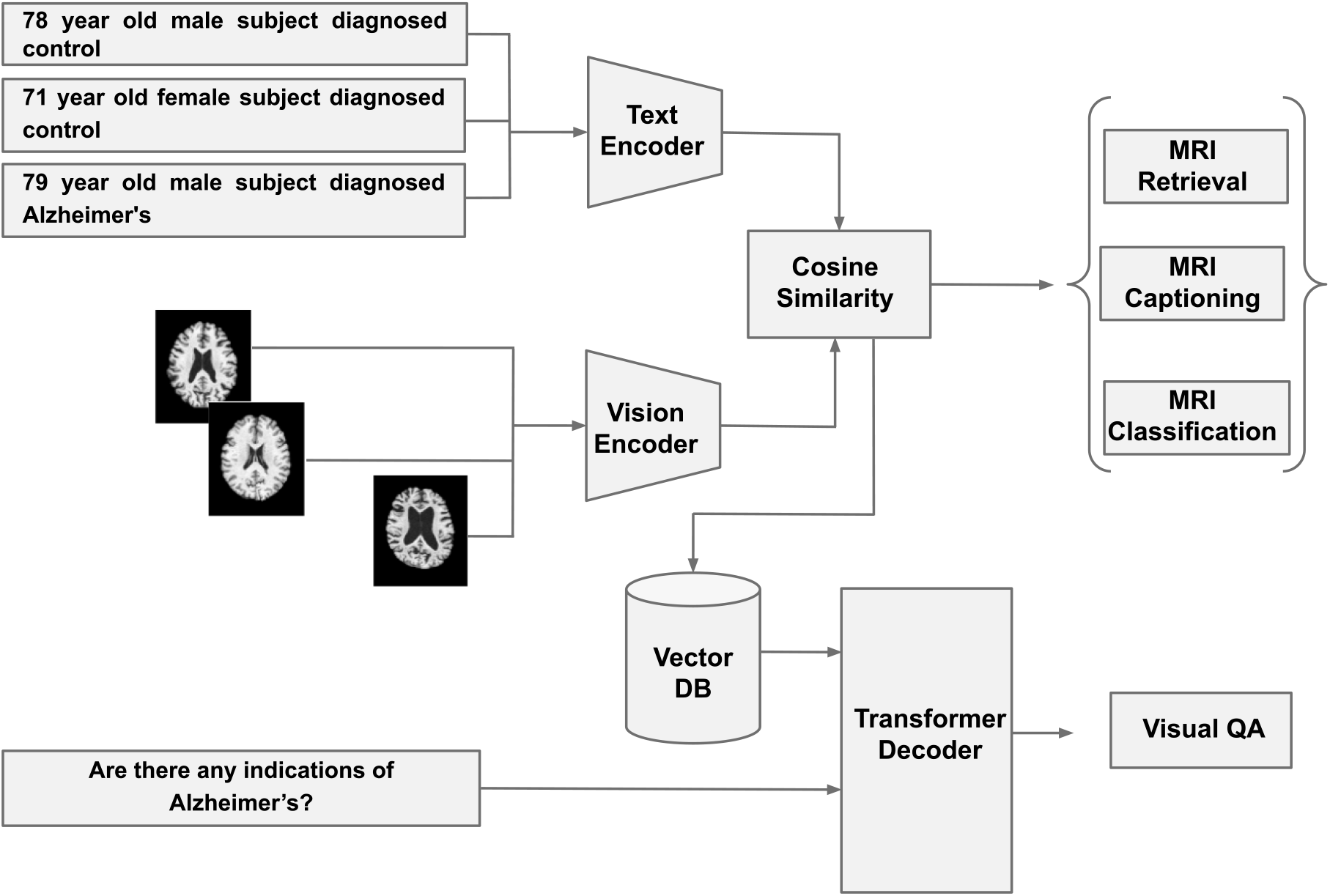
The Proposed Vision-Language Model. The left-hand side of the figure shows the possible user-side inputs (clinical text only, image only, image + clinical text, image + clinical text + user query). The right-hand side shows the response from the model (MRI retrieval, MRI captioning, MRI classification, visual question-answering).

## II. Data

In this study, we used the widely-used and publicly available ADNI (Alzheimer’s Disease Neuroimaging Initiative) brain MRI datasets for our analyses [18] (**Table 1**). We pre-processed the 3D T1-weighted MRI scans using a standard set of pre-processing steps [19] including: nonparametric intensity normalization (N4 bias field correction) [20], ‘skull-stripping’, linear registration to a template with 9 degrees of freedom, and isotropic resampling of voxels to 2-mm resolution. The input dimension of the MRI spatially was 91×109×91. All images in the dataset was z-scored for standardization.

**TABLE I.**
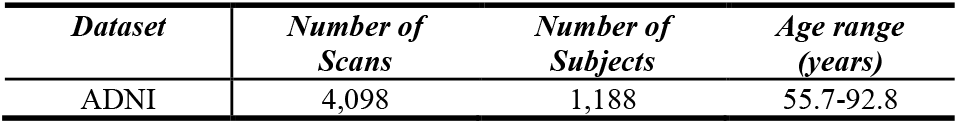
Volumetric T1-weighted Brain MRI dataset.

## III. Methods

### Stage 1: Uni-Modal Pre-training

For our tasks, we want to be able to input text—either in the form of text prompts or queries, or as sentences describing the imaging data in text format—so their relationship can be learned. We tested several transformer-based text encoders to extract strong embeddings from the text inputs. This included BERT [21], ClinicalBioBERT [22], DeBERTa [23], BiomedBERT [24], and Sentence BERT [25]. We also created and evaluated different prompt structures to incorporate the covariates. Our final prompt format followed the style “78 year old male subject diagnosed [with] Alzheimer’s”. We selected the Sentence BERT as our main text encoder given its strong out-of-the-box performance on numerical and categorical concepts. Since “Alzheimer’s” was not already a part of the model’s vocabulary (the collection of words that the model can understand or generate), it was added to the tokenizer as a new token (in fact, we used the token “alzheimers” to standardize tokenization). We conducted domain-specific pretraining on the text encoder using masked language modeling [21] (MLM). We created a synthetic pre-training dataset of 25,000 text captions varying the age, sex and the disease descriptor. Here we masked 35% of the input text tokens and pre-trained the model to predict and thereby improve understanding of the numerical and categorical variables.

Further, we used the powerful 3D DenseNet121 CNN [26] as our image encoder. This architecture has been proven [9] to be effective in extracting discriminative features for Alzheimer’s disease. The image encoder was pre-trained using contrastive learning and fully supervised learning with T1-weighted MRIs from ADNI [9]. The outputs of both the image and text encoder are passed through individual projection heads.

### Stage 2: Cross-Modal Contrastive Language-Image Fine-tuning

Given a batch of *N* image-text pairs {(*I*_*i*_, *T*_*i*_)}^N^_i=1_, the similarity matrix s_i,j_ is defined as in equation (1), as the L2-normalized dot product between each image and text embedding:

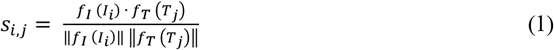

Here, *f*_*I*_ and *f*_*T*_ are the vision and text encoders; *I*_*i*_ and *T*_*j*_ are a given image and text pair.

The symmetric cross-entropy loss ℒ is defined as in equation (2):

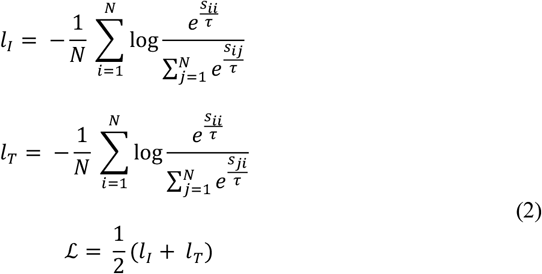

Here, *l*_*i*_ and *l*_*T*_ are the image-to-text and text-to-image losses, and ***τ*** is the temperature parameter that controls the range of the logits – this parameter is directly optimized during training, as a log-parameterized multiplicative scalar.

The pre-trained vision-language models were fine-tuned using different methods - full fine-tuning of all parameters; partial fine-tuning of all parameters after and including the last two layers of the encoders; locked image tuning [27] - freezing the image encoder backbone and fine-tuning everything else; or fine-tuning only the two projection heads.

### Stage 3: Applications

The first application we trained the model to perform was MRI retrieval, i.e., given a text prompt, the framework retrieves the top-*k* image results corresponding to the text. Conversely, MRI captioning involves retrieving or generating the top-*k* text caption results given an input image. For MRI image classification, given an image and covariates (included as encoded text; see below), the model outputs a binary classification result that the patient has Alzheimer’s disease or is a healthy control. We also tested the effect of re-ranking on the retrieval results using a cross-encoder approach [28] [29].

We evaluated multiple state-of-the-art contemporary transformer-based decoder-only large language models (LLMs) for visual question answering including Google’s Gemma2 series [30], Meta’s Llama3 series [31], and Mistral’s 7B [32]. The Mistral 7B LLM was selected due to its strong ability to adhere to provided instructions. We used Meta AI’s optimized FAISS vector store [33] to create our database of vector embeddings for retrieval and re-ranking mechanisms for visual question answering. The retrieved results were re-ranked based on the corresponding covariates provided as an additional input.

## IV. Experiments AND Results

### A. Setup

We performed a random search to select hyperparameter values, including the learning rate {1e-3 to 1e-6}, weight decay between 0.01 and 0.0001, batch size {4, 8, 16, 32, 64, 128, 256}, projection head dimension {256, 512, 1024}. The ADAM optimizer was used in all experiments. For the retrieval experiments, *k* was set to 10. A cosine learning rate scheduler with warmups was used for the image and text encoders, as well their respective projection heads.

For our retrieval tasks (text-to-image and image-to-text) we used three standard evaluation metrics that are widely used in machine translation - specifically, bilingual evaluation understudy (BLEU), Recall-Oriented Understudy for Gisting Evaluation - Longest Common Subsequence (ROUGE-L)-- both to evaluate the quality of overlap between the retrieved result and the ground truth–as well BERTScore to quantify the semantic similarity between the two based on token embeddings. In addition, we used the receiver-operator characteristic curve-area under the curve (ROC-AUC), Accuracy for classification and Mean Absolute Error (MAE) for age prediction. The retrieval results and the VQA were calculated using over 100 unique retrievals. The classification results were calculated over 1,219 T1-weighted MRI scans.

We performed a series of ablation experiments, including ablations within the text encoder, the text pre-training, the image encoder pre-training, the cross-modal fine-tuning method. These experiments are summarized in the following results sections per application.

### B. Application 1: MRI Retrieval

We tested the fine-tuned cross-modal model for MRI retrieval using unique text captions sampled from the test set as the inputs. The results of these tests are presented in **Table 2** as an average using the top-10 retrieved images.

**TABLE II.**
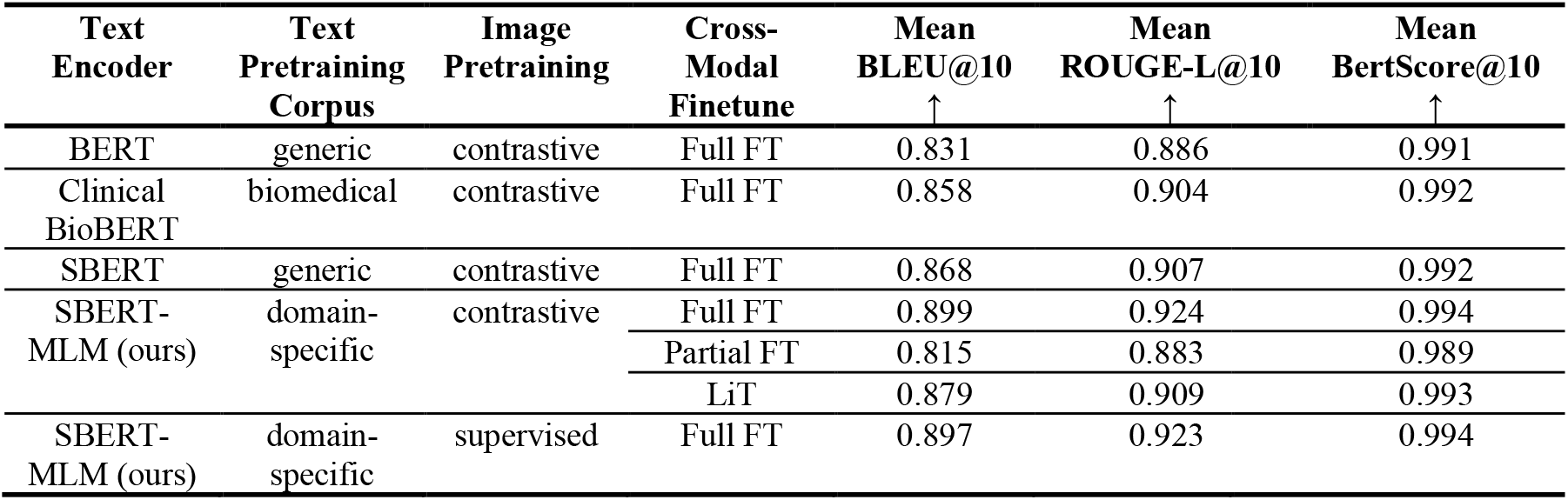
MRI retrieval results (Text-to-Image), where FT is fine-tune, LIT is locked image tuning, MLM is masked language modeling.

### C. Application 2: MRI Captioning

We tested the fine-tuned, cross-modal model for MRI captioning, with unique images sampled from the test set as inputs. The results of these tests are summarized in **Table 3** as an average using the top-10 retrieved text captions.

**TABLE III.** MRI Captioning results (Image-to-Text).

| Text Encoder | Text Pretraining Corpus | Image Pretraining | Cross-Modal Finetune | Mean BLEU@10<br>↑ | Mean ROUGE-L@10<br>↑ | Mean BertScore@10<br>↑ |
| --- | --- | --- | --- | --- | --- | --- |
| BERT | generic | contrastive | Full FT | 0.544 | 0.796 | 0.953 |
| Clinical BioBERT | biomedical | contrastive | Full FT | 0.512 | 0.787 | 0.953 |
| SBERT | generic | contrastive | Full FT | 0.500 | 0.783 | 0.950 |
| SBERT-MLM (ours) | domain-specific | contrastive | Full FT | 0.579 | 0.803 | 0.955 |
|  |  |  | Partial FT | 0.422 | 0.760 | 0.939 |
|  |  |  | LiT | 0.571 | 0.799 | 0.957 |
| SBERT-MLM (ours) | domain-specific | supervised | Full FT | 0.654 | 0.821 | 0.957 |

### D. Application 3: MRI classification

We tested the fine-tuned cross-modal model for MRI classification with unique images sampled from the test set along with their covariates. Results of these tests are presented in **Table 4** as an average using the top-10 retrieved text captions along with ablation runs.

**TABLE IV.** MRI Classification results.

| Text Encoder | Text Pretraining Corpus | Image Pretraining | Cross-Modal Finetune | Mean ROC-AUC<br>↑ |
| --- | --- | --- | --- | --- |
| BERT | generic | contrastive | Full FT | 0.683 |
| Clinical BioBERT | biomedical | contrastive | Full FT | 0.689 |
| SBERT | generic | contrastive | Full FT | 0.709 |
| SBERT-MLM (ours) | domain-specific | contrastive | Full FT | 0.786 |
|  |  |  | Partial FT | 0.768 |
|  |  |  | LiT | 0.778 |
| SBERT-MLM (ours) | domain-specific | supervised | Full FT | 0.827 |

### E. Application 4: Visual Question Answering

A sample VQA interaction with our model is shown in **Table 5**. Here the model was given an image along with an instruction and query. Additionally, covariates are provided as encoded text to the model and used to improve the retrieval performance via re-ranking. The responses from the model are used to quantify its performance using average MAE (in years) for age prediction and average accuracy for disease classification, over the test set of T1-weighted MRIs.

**TABLE V.** Visual Question Answering results.

| Text Encoder | Text Pretraining Corpus | Image Pretraining | Cross-Modal Finetune | AD Classification Mean Accuracy<br>↑ | Age Prediction Mean MAE<br>↓ |
| --- | --- | --- | --- | --- | --- |
| SBERT-MLM (ours) | domain-specific | contrastive | Full FT | 0.780 | 4.820 |
| SBERT-MLM (ours) | domain-specific | supervised | Full FT | 0.810 | 5.570 |

**TABLE VI.**
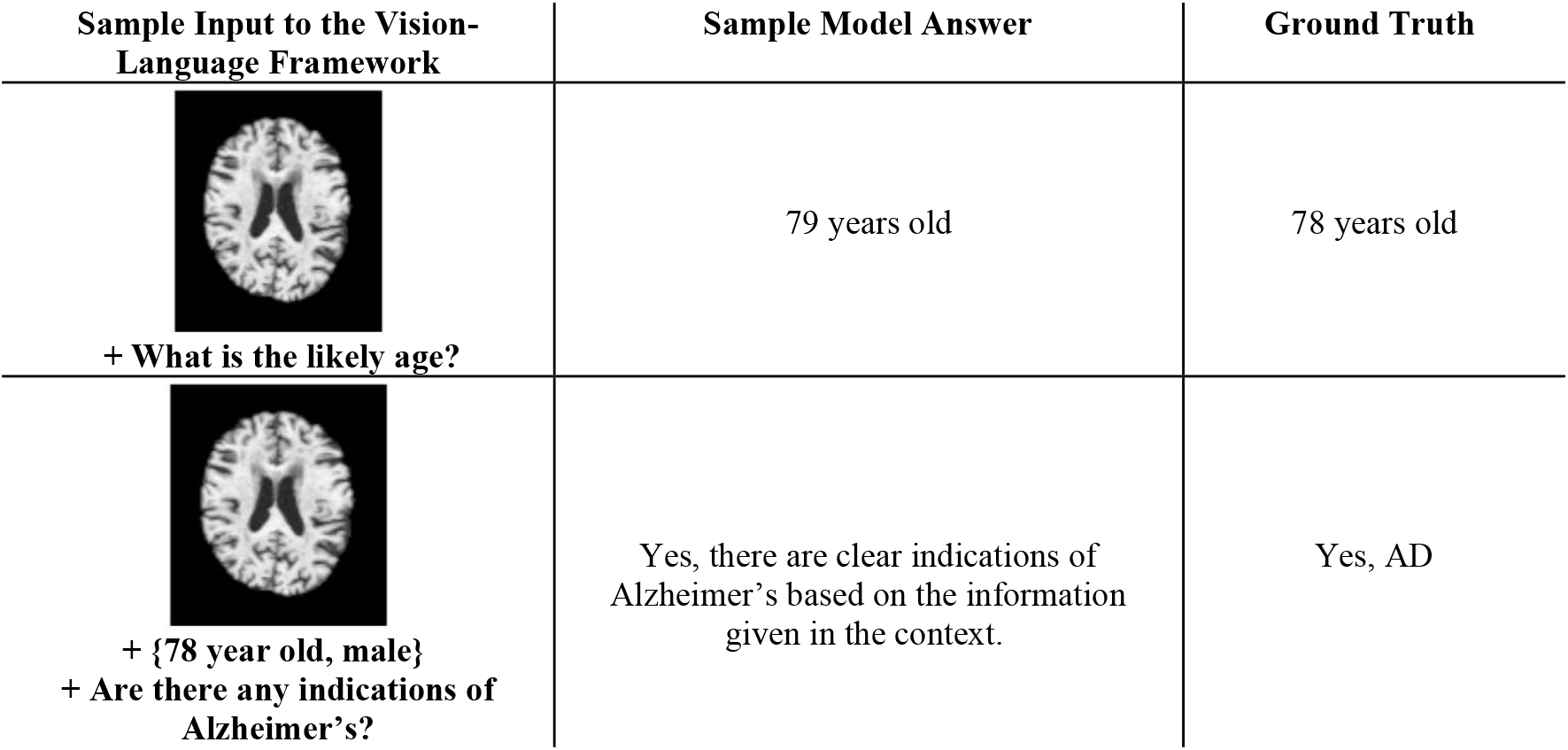
Sample VQA interaction with the proposed model, given that the ground truth is: 78 year old male diagnosed with Alzheimer’s disease.

| Sample Input to the Vision-Language Framework | Sample Model Answer | Ground Truth |
| --- | --- | --- |
| + What is the likely age? | 79 years old | 78 years old |
| + {78 year old, male}<br>+ Are there any indications of Alzheimer’s? | Yes, there are clear indications of Alzheimer’s based on the information given in the context. | Yes, AD |

## V. Discussion

### A. The pretraining and encoder architectures provide critical initialization for cross-modal retrieval tasks

In our experiments, the text encoder and its pre-training corpus played a key role in cross-modal retrieval and classification performance, as seen in **Tables 2, 3**, and **4**. We observed that most ‘off-the-shelf’ text encoders were not sensitive to numerical and categorical concepts that are crucial to neuroimaging data. We also found that the cross-modal tasks were influenced by the image encoder’s pre-training. Specifically, while supervised pre-training improved downstream classification of the same task, contrastive pre-training provided a more balanced overall performance across all applications tested.

### B. Cross-modal fine-tuning methods can be more parameter-efficient but still maintain performance

We evaluated different cross-modal fine-tuning methods - even though fully fine-tuning all layers usually yielded the best performance, the locked image tuning – which fine-tuned only the text backbone along with the image projection head – had competitive performance. This greatly reduces the total number of tunable parameters, and lends itself to low-resource settings.

### C. Strong vector embeddings lead to strong downstream results

We created a vector database from our trained cross-modal model to facilitate visual question answering by extending the capabilities of an instruction-tuned transformer decoder model.

The VQA performance on key benchmarking tasks such age prediction and disease detection were competitive with stand-alone regression and classification models. This shows that a single cross-modal model can handle multiple tasks without any additional fine-tuning. It is perhaps surprising that strong predictive models of age and disease are implicitly learned from the text-image embeddings, whereas usually classifiers are custom-trained using labeled training data, where the labels are typically a single parameter (age or diagnosis).

Future work will include additional evaluation of our vision-language model on other datasets, modalities, and disorders.

## VI. Conclusion

In this work, we presented a novel vector retrieval-driven vision-language framework that enhances brain image analysis by integrating T1-weighted imaging with complementary non-imaging clinical data. Although the approach is quite general and would be applicable to a variety of applications in radiology and medical imaging, we illustrated the approach on several tasks that arise in Alzheimer’s disease research. In the future, using an LLM interface, such tasks could be composed to perform virtual experiments, such as identifying brain abnormalities in a group of patients, or discovering imaging patterns associated with a medication or risk factor described in the associated text. Our approach leverages cross-modal retrieval techniques and fine-tuning of multimodal embeddings to enhance diagnostic assessments, providing a framework to capture subtle disease-related patterns that may be overlooked by single-modality analyses.

Overall, this study offers a promising step toward more robust, data-driven approaches in radiologic research, with illustrative examples highlighting the role of cross-modal analysis in advancing our understanding of neurodegenerative diseases.

## Notes

This work was supported by the U.S. National Institutes of Health, under NIH grant U01 AG068057 (‘AI4AD’).

### Competing Interest Statement

The authors have declared no competing interest.

